# Acoustic prediction and biomechanical validation of primary stability in uncemented short-stem hip prostheses: an experimental study

**DOI:** 10.1101/2025.05.28.656119

**Authors:** Alexander Jahnke, Simon Schreynemackers, Ahmed Tawous, Swantje Petersen, Samar Hamad, Markus Rickert, Bernd Ishaque

## Abstract

**Background:** In uncemented hip arthroplasty, achieving sufficient primary stability is essential for long-term implant success. However, objective intraoperative assessment of fixation quality remains challenging. Acoustic analysis of stem impaction sounds offers a promising tool for real-time evaluation, but its diagnostic accuracy and biomechanical correlation require further validation.

**Methods:** Twelve formalin-fixed human femora were implanted with cementless Metha® short stems under three predefined anchorage conditions: loose, optimal (fit), and fracture-inducing press-fit. Impaction sounds were recorded using calibrated microphones and processed via frequency-domain analysis. Relative micromotions were quantified under torsional loading to biomechanically assess primary stability.

**Results:** Spectral markers reliably differentiated between anchorage states. The transition from loose to fit showed minimal spectral change, while fit-to-fracture was characterized by a significant increase in low-frequency energy (<2.5 kHz) and pronounced attenuation in high-frequency bands (>15 kHz). These acoustic signatures closely correlated with biomechanically measured micromotions, which showed a distinct hierarchy: fracture < fit < loose. Cluster permutation analysis confirmed statistically significant differences, particularly in the fracture group.

**Conclusion:** This in vitro study demonstrates that frequency-based acoustic analysis can distinguish between stable, insufficient, and over-press-fit conditions during stem implantation. The findings support the potential of intraoperative acoustic monitoring as a real-time, objective tool to enhance implant safety and detect cortical compromise before it becomes clinically apparent.

## Introduction

The implantation of hip endoprostheses is one of the most common surgical procedures in orthopaedic surgery and in many cases leads to a significant improvement in quality of life (Grubor et al. 2013). Despite established surgical techniques and the use of high-quality implant systems, complications such as loosening, periprosthetic fractures or the need for revision surgery remain a relevant clinical problem. Intraoperative fractures in particular occur with a frequency of around 1-5% (Kenanidis et al. 2020). With uncemented implants, this risk is particularly relevant due to the necessary interference fit, which results from an undersized prosthesis bearing in the bone and into which the prosthesis is driven (Hailer et al. 2010).

Postoperative primary stability, i.e. the direct mechanical anchoring of the prosthesis in the bone, is crucial for the success of the treatment. It is the prerequisite for subsequent bony integration and thus for the long-term stability of the implant (Bürkner et al. 2012). However, there is currently no reliable, commercially available system for the objective intraoperative assessment of this stability. Surgeons are currently highly dependent on their experience and subjective perception during implantation - for example through tactile feedback when the prosthesis is inserted. Whether this results in fine fissures or even fractures can often not be determined with certainty intraoperatively. Even imaging procedures such as intraoperative X-ray checks offer only limited certainty here, as possible fractures can easily be overlooked due to the overlapping of the prosthesis in the bone (Hassan et al. 1998; HARRIS et al. 1991).

Promising approaches to improving intraoperative assessment are currently focusing on analyzing acoustic signals that are generated during implantation. These change depending on the anchoring of the implant in the bone and allow conclusions to be drawn about the strength of the connection and possible microfractures (García-Vilana et al. 2025; Aggelis et al. 2015; Sakai et al. 2011). In a previous study, it was shown that the stability of the femoral stem can be reliably monitored in real time using acoustic methods. The results underline the importance of sufficient primary stability for successful bony integration and the long-term function of the implant (Fonseca Ulloa et al. 2023). Acoustic methods could therefore make an important contribution to optimizing surgical results by providing immediate feedback.

This in vitro study aims to determine whether frequency-domain analysis of impaction sounds can serve as a reliable, objective predictor of primary stability in uncemented short-stem hip prostheses. Furthermore, it seeks to validate the diagnostic value of acoustic signatures by correlating them with biomechanically quantified micromotions under controlled torsional loading.

## Material and Methods

### Human specimen

A total of n=12 formalin-fixed human femora (nine right, three left) were used for the present study. The specimens came from body donors of the Anatomical Institute of the Justus Liebig University Giessen. Ethical approval for the use of the specimens has been granted under file number 160/19.

### Prostheses

The cementless Metha^®^ short stem prostheses (B. Braun, Aesculap AG, Tuttlingen, Germany) were used. Prosthesis sizes 4 to 7 with a caput-collum-diaphyseal angle of 120° were used.

### Preparation and planning

After harvesting, the femur specimens were largely freed of soft tissue and radiologically measured in the anterior-posterior beam path to determine the appropriate prosthesis size. The collum was then prepared according to the radiological planning using the MediCAD software (mediCAD Hectec GmbH, Altdorf /Landshut, Germany). The medullary cavity was rasped to the respective size using the manufacturer’s instruments. The bones were numbered and deep-frozen at -20°C until the day of implantation.

### Implantation of the prostheses

Before implantation, the bones were thawed at 16 ± 1°C room temperature for 3 hours. The distal end of the prostheses was drilled in advance with a 1.9 mm drill for the subsequent application of a measuring pin for primary stability measurement. To prevent excessive movement during implantation, the femora were fixed in the desired position in a non-slip hard foam block (PE block foam, Wilhelm Julius Teufel GmbH, Germany). This was then pressed against the laboratory operating table using two clamps and an assistant (A. T.). The implantation itself was performed by an experienced specialist (BA. I.). The specimens were randomly assigned to three different study groups with n=4 specimens each: too loose (*loose*), correct (*fit* - which refers to the surgeon’s intraoperative assessment of optimal press-fit without visible cortical damage) and up to fracture of the bone (*fracture*). These conditions were documented and recorded by the surgeon. A detailed overview of the femoral specimens and prosthesis sizes in the different groups is shown in Table 1 (Table 1).

**Table 1:** Status assignment of the femora (R = right, L = left) and prosthesis sizes.

| State | Specimen | Prosthesis size |
| --- | --- | --- |
| fit | #13 R | 4 |
| fit | #13 L | 4 |
| fit | #3 R | 6 |
| fit | #9 R | 7 |
| loose | #5 L | 4 |
| loose | #4 R | 4 |
| loose | #14 L | 5 |
| loose | #14 R | 6 |
| fracture | #3 L | 4 |
| fracture | #10 R | 5 |
| fracture | #5 R | 6 |
| fracture | #9 L | 7 |

### Acoustic instrumentation

Impaction sounds were recorded with a laboratory-calibrated measurement microphone (MM 1, beyerdynamic, Heilbronn, Germany; free-field response 20 Hz – 20 kHz, ±1.5 dB). The capsule was positioned 50 cm from the prosthesis axis, aimed directly at the stem shoulder. The microphone fed a four-channel USB interface (Rubix 44, Roland Corporation, Hamamatsu, Japan; 24-bit, 192 kHz, ASIO), which served as external A/D converter. Prior to the first implantation a calibration blow was used to set the analogue gain such that the loudest strike remained ≥6 dB below full scale; the setting was kept unchanged for all specimens. Signals were stored as 44.1 kHz/24-bit PCM.

### Signal processing

Individual hammer blows were isolated by a back-tracked spectral flux onset detector implemented in *librosa 0.10.1.* Each segment was zero-padded or truncated and transformed with a single-sided FFT of equal length. The amplitude spectrum P_l_ was energy-normalised 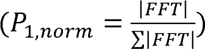 to compensate for amplitude variability; segments whose total energy fell below five percent of the median were discarded. Four regions of interest (ROI) reproduced the “original bands” reported by Fonseca et al. (Fonseca Ulloa et al. 2023): Low (0–2.5 kHz), 2.9 kHz (2.7–3.1 kHz), 4.4 kHz (4.2–4.6 kHz) and 8.7 kHz (8.5–8.9 kHz).

### Software environment

All signal processing and statistical procedures were executed in Python 3.9 under Anaconda (Win-64). Core numerical operations relied on NumPy 1.26.1 and SciPy 1.13.1, whereas audio segmentation and resampling were handled by librosa 0.10.1 and soundfile 0.12.1. The non□parametric statistics were implemented with MNE-Python 1.8.0 (cluster permutation) and Pingouin 0.5.5 (permutation *t*, effect sizes). Data management used pandas 2.1.1; graphics were created with Matplotlib 3.8.0. Machine-learning benchmarks employed scikit-learn 1.3.1. Low-level FFT kernels were accelerated by Numba 0.58.1.

### Preparation of the primary stability measurement

For the primary stability measurement, the femora were then embedded in epoxy resin (GP 010 A/B, Gößl + Pfaff GmbH, Germany) at the condyles. A reference frame was then fixed at the level of the lesser trochanter. This served as a base for attaching the measuring frame with integrated measuring sensors and as a reference point. Starting from this reference point, the positions of the bone measuring points could be precisely determined. The bone measuring points were 45 mm proximal (B_1_) and 30 mm (B_2_) and 80 mm (B_3_) distal to the lesser trochanter. Holes with a diameter of 1.9 mm were drilled at each of the previously defined measuring points and the measuring pins were fixed in place using superglue. In addition, another pin was attached to the proximal shoulder of the prosthesis (P_p_). At the distal end of the prosthesis, the surrounding bone was specifically expanded with a 10 mm drill to expose the hole previously made. The distal measuring point (P_d_) could then be precisely inserted and fixed in this hole.

### Primary stability measurement

Based on previous studies, retrograde and reaction-free axial torsional moments around the longitudinal Z-axis of the femoral stem were applied into the prosthesis. These torques were increased in a continuous process up to a maximum value of ±3.5 Nm (for *fit* and *loose*) and ±1.75 Nm (for *fracture*) in 80 incremental steps. The maximum torque was reached twice in each direction of rotation, resulting in 480 individual measurements per series. Each series of measurements was repeated three times for each measuring point (Fölsch et al. 2024; Jahnke et al. 2020; Fölsch et al. 2023).

This method was used to generate very small but measurable micromovements, if possible without impairing the respective anchoring stability of the prosthesis. To record these movements, the measuring pins at the bone measuring points (B_1_-B_3_) as well as at the prosthesis (P_p_ + P_d_) were connected to the individual measuring points one after the other via a measuring cube. This cube served as a local coordinate system. After each torsion step, the spatial position of the measuring cube was recorded using 9 non-contact eddy current sensors (NCDT 3010-S2, Micro-Epsilon Messtechnik GmbH & Co. KG, Ortenburg, Germany) in a 3-3-3 arrangement per plane with a resolution of 0.1 μm. The normalized rotational stability was then calculated for each measurement point as a function of the applied torque (mdeg/Nm) in order to characterize the anchoring behaviour of the prosthesis. The proximal and distal relative micromovements (rm_1_ and rm_2_) were calculated as the difference between the corresponding measuring point heights of the prosthesis and bone measuring points P_p_-B_1_ (rm_1_) and P_d_ - B_2_ (rm_2_) (Figure 1).

**Figure 1:**
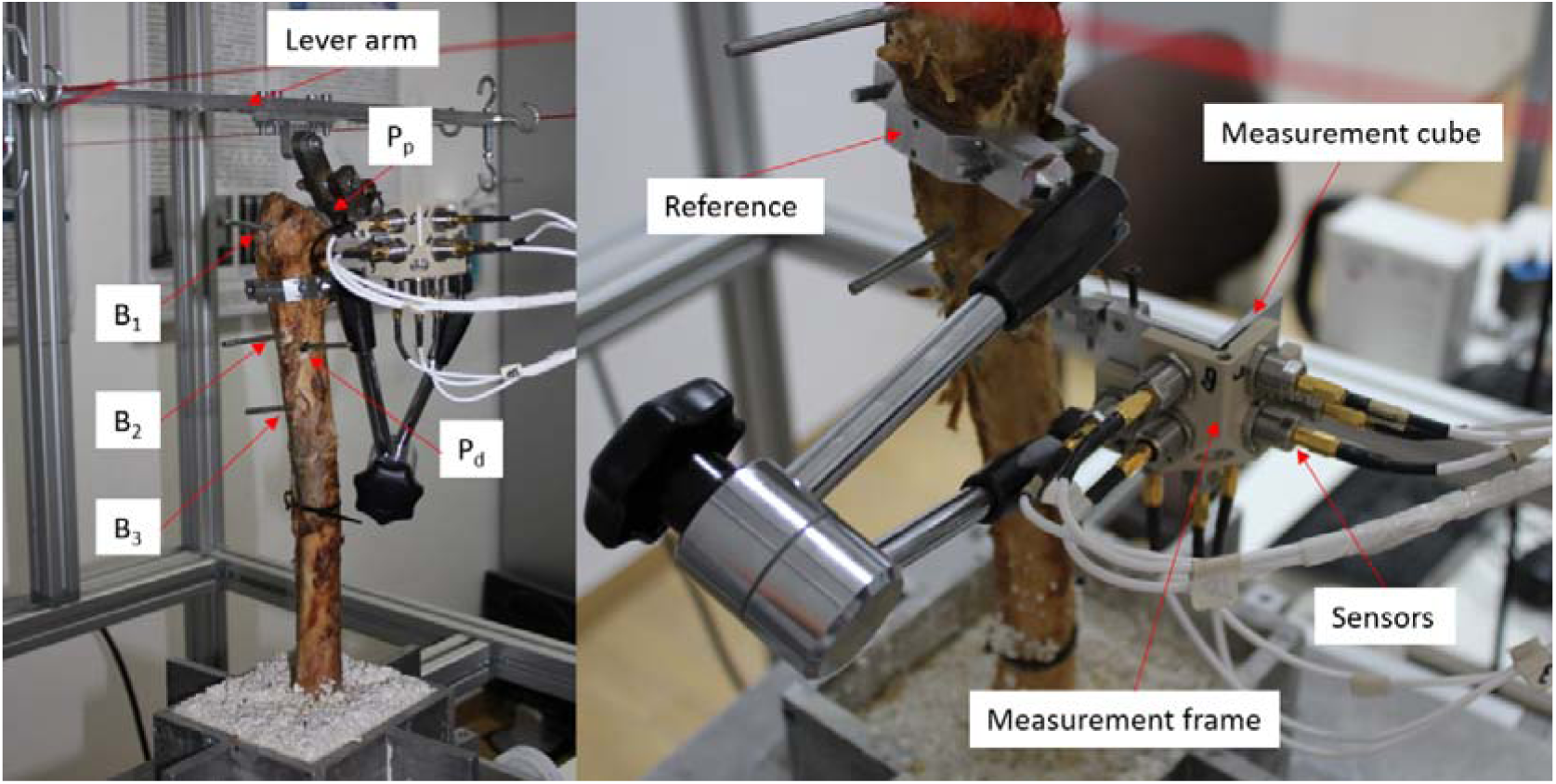
Experimental setup and measurement protocol for quantifying primary stability via torsion-induced micromovements

Due to the different lengths of the prosthesis models used, the relative movements rm_1_ and rm_2_ were transferred to the level of the corresponding bone measuring points B_1_ and B_2_ using linear interpolation. This ensures a methodologically consistent and statistically comparable evaluation of the different stem designs (Jahnke et al. 2020).

### Statistics

Band-wise differences (Δµ) between conditions were assessed with independent permutation *t*-tests (10 000 resamples, two-tailed); effect size was expressed as Cohen’s *d*. Whole-spectrum differences were quantified by the two-sample Kolmogorov–Smirnov statistic. Frequency-resolved effects were detected with a one-dimensional cluster permutation test (5 000 permutations, α = 0.05). An adjacency matrix connected frequency bins within a gap of ≤20 bins, preventing fragmentation; clusters with family-wise *p* < 0.05 were deemed significant. The statistical evaluation of the measurement results of the primary stability analysis was carried out using SPSS software (version 29.0, IBM, New York, United States).

A non-paremetric Kruskal-Wallis test was performed to identify significant differences between the groups. The results were evaluated at a significance level of p < 0.05.

## Results

### Band-wise permutation analysis

Table 2 summarises the permutation-based comparisons (10 000 resamples, two-tailed) within the four frequency windows originally reported by Fonseca et al. (Fonseca Ulloa et al. 2023). Effect sizes are expressed as Cohen’s *d*. Only the fit–fracture and loose–fracture contrasts revealed statistically relevant differences (α<0.05, family-wise), predominantly in the low-frequency band (< 2.5 kHz) and the first high-frequency window around 8.7 kHz. In contrast, loose–fit differences were negligible (|d| ≤ 0.14, *p*>0.3).

**Table 2:** Band-wise permutation analysis reveals significant fit–fracture and loose–fracture differences in selected frequency bands.

| original band | Comparison | $\Delta\mu$ | $p$-value | $d$ | Interpretation |
| --- | --- | --- | --- | --- | --- |
| 0–2 500 Hz | loose–fit | $5.5 \times 10^{-4}$ | 0.967 | +0.01 | no effect |
| | fit–fracture | **+0.062** | **0.0002** | +1.02 | substantial ↑ in fracture |
|  | loose–fracture | +0.063 | 0.007 | +0.58 | moderate |
| 2 700–3 100 Hz | loose–fit | +0.0020 | 0.38 | +0.14 | trivial |
|  | fit–fracture | **+0.0069** | **0.034** | +0.46 | medium |
|  | loose–fracture | **+0.0089** | **0.0006** | +0.76 | large |
| 4 200–4 600 Hz | loose–fit | –0.00027 | 0.86 | –0.03 | none |
|  | fit–fracture | +0.0014 | 0.506 | +0.16 | small |
|  | loose–fracture | +0.0011 | 0.578 | +0.12 | small |
| 8 500–8 900 Hz | loose–fit | +0.00052 | 0.515 | +0.10 | none |
|  | fit–fracture | **−0.0040** | **0.0002** | −1.37 | large ↓ in<br>fracture |
|  | loose–fracture | **−0.0035** | **0.011** | −0.62 | moderate |

### Global distribution tests

Kolmogorov–Smirnov tests, performed on the complete spectra (Table 3), confirmed global distributional differences between all three conditions. The largest separations were observed for pairings that involved the fracture state.

**Table 3:**
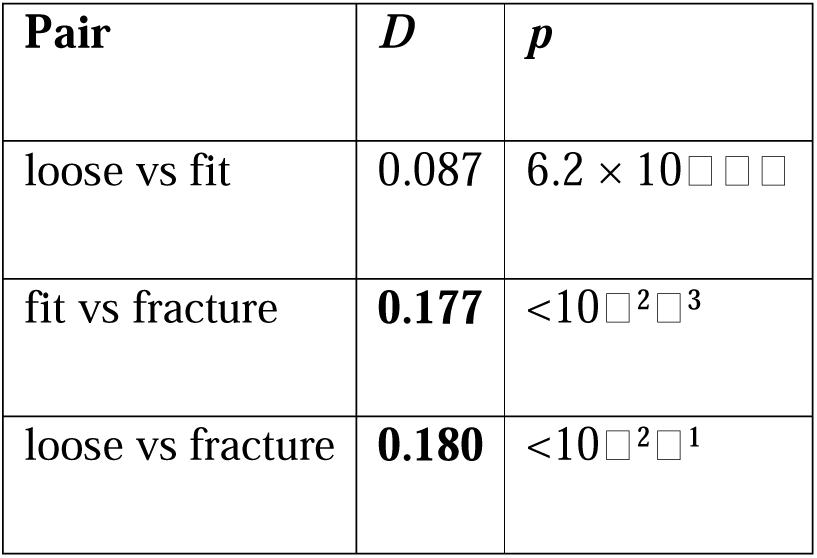
Kolmogorov–Smirnov tests confirm global spectral differences, with largest deviations involving the fracture condition.

### Cluster permutation tests

A one-dimensional cluster permutation test (5 000 permutations; threshold *t*-value corresponding to α = 0.05; maximum spatial gap ≤ 20 frequency bins) was applied to the log-spectral amplitudes. Nine, 16 and 12 significant clusters (family-wise α < 0.05) were obtained for loose–fit, fit–fracture and loose–fracture respectively (Figure 4–6).

### Visual summary

Stacked plots (Figure 2: loose–fit, Figure 3: fit–fracture, Figure 4: loose–fracture) display mean ± SD spectra with super-imposed cluster spans and the absolute *T* metrics. Legends identify every significant cluster by its rounded 100-Hz limits and corrected *p*-value.

**Figure 2:**
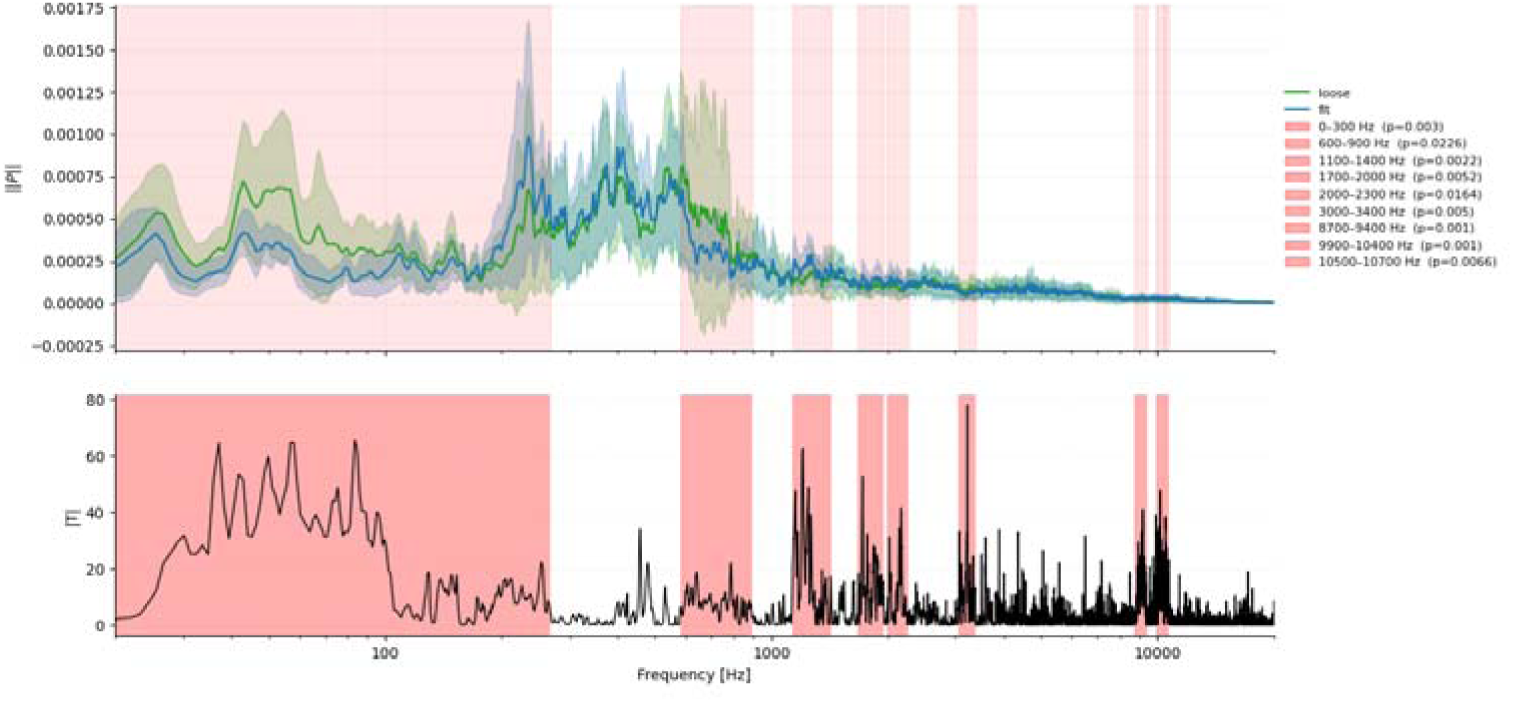
**Loose vs fit**: the original bands reported by Fonseca et al. (Fonseca Ulloa et al. 2023) (< 2.5 kHz, 2.9 kHz, 4.4 kHz, 8.7 kHz) were all replicated, although their summed |*T*| ranked only second to eighth (e.g., 8 724–9 380 Hz, *p*=0.001).

**Figure 3:**
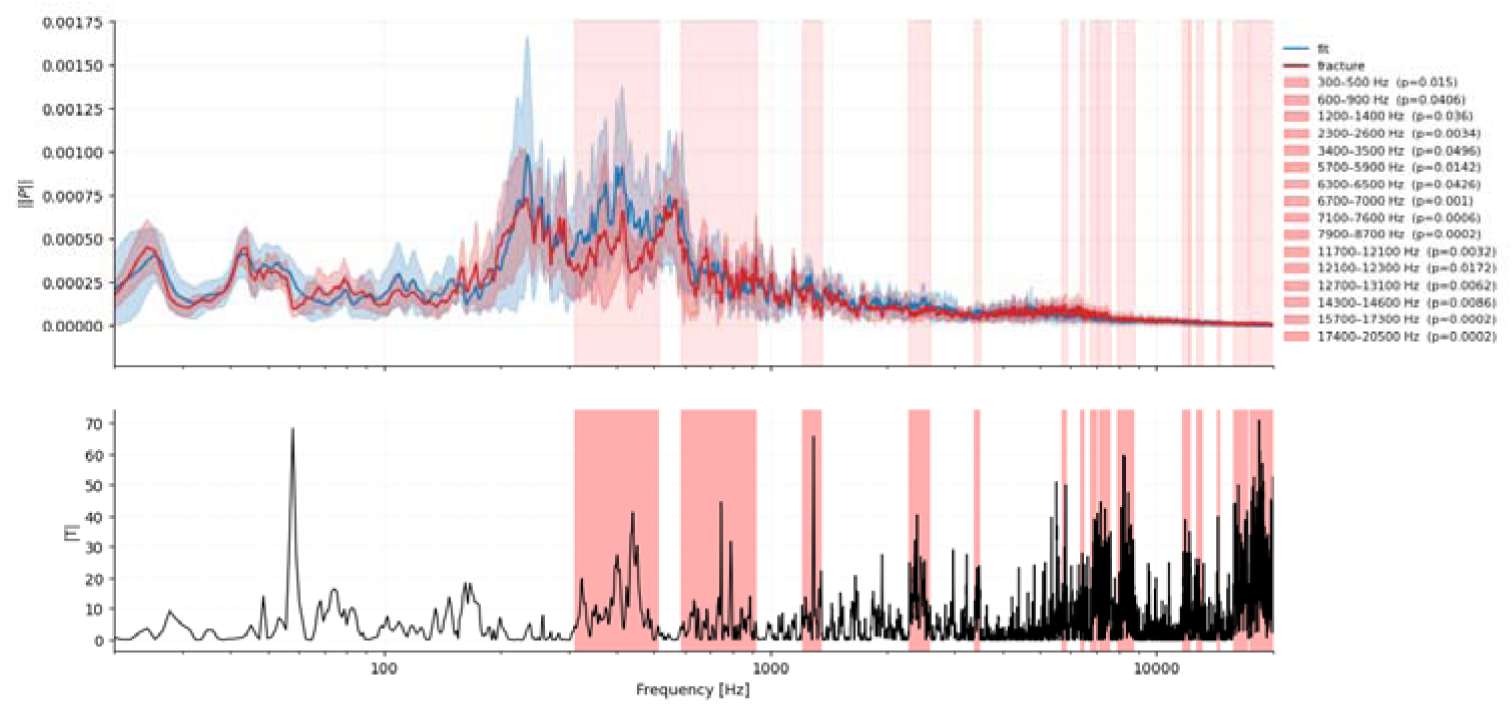
**Fit vs fracture**: new, wide clusters emerged between 15–20 kHz and 7–9 kHz (Σ|*T*| > 22 000; *p*≤0.0002), whereas the classical bands shifted upwards by ≈ 0.5 kHz.

**Figure 4:**
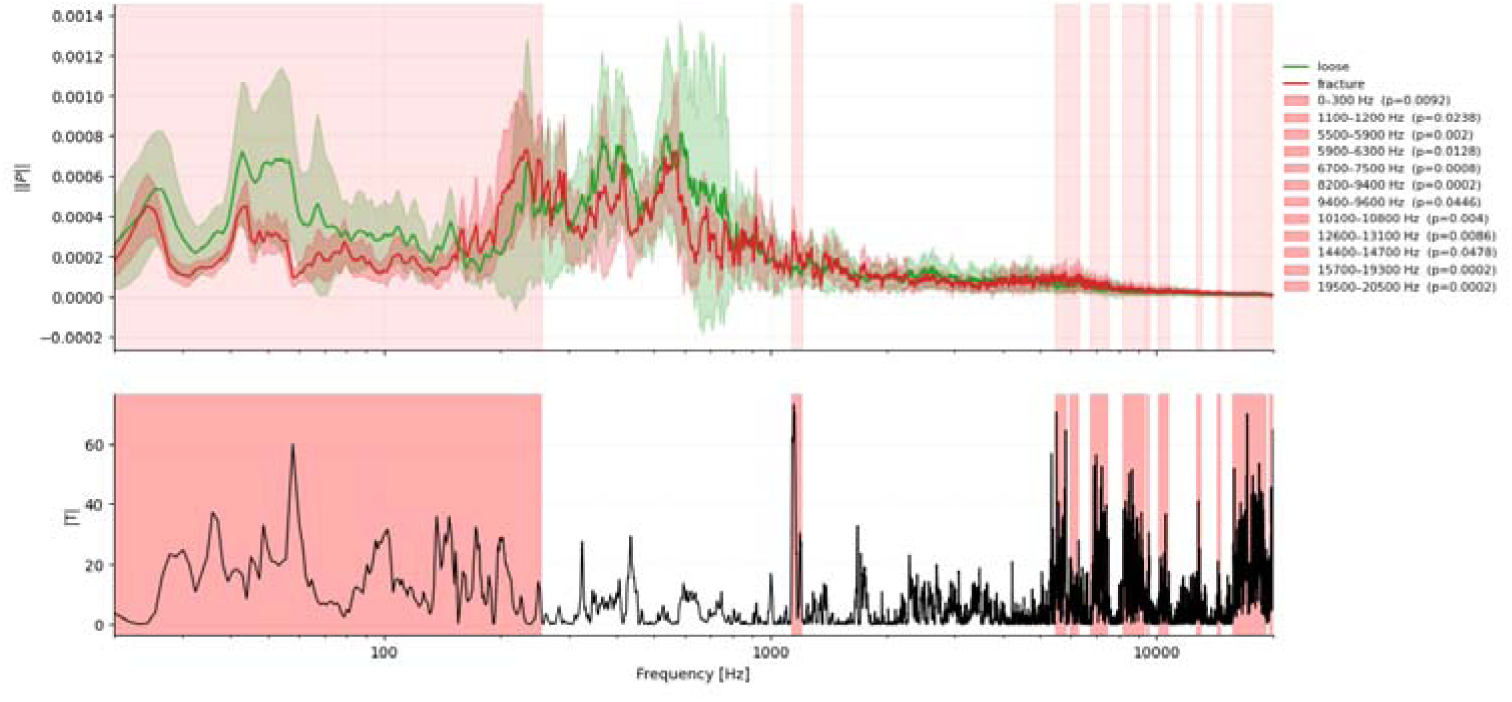
**Loose vs fracture**: combined effects yielded two dominant high-frequency clusters (15–19 kHz, 8–9 kHz) and a broad low-frequency cluster (0–255 Hz, *p*=0.009).

A representative image of an intraoperative fracture is shown in Figure 5. The cortical disruption (indicated by the red arrow) occurred during stem impaction and reflects a critical outcome of excessive press-fit.

**Figure 5:**
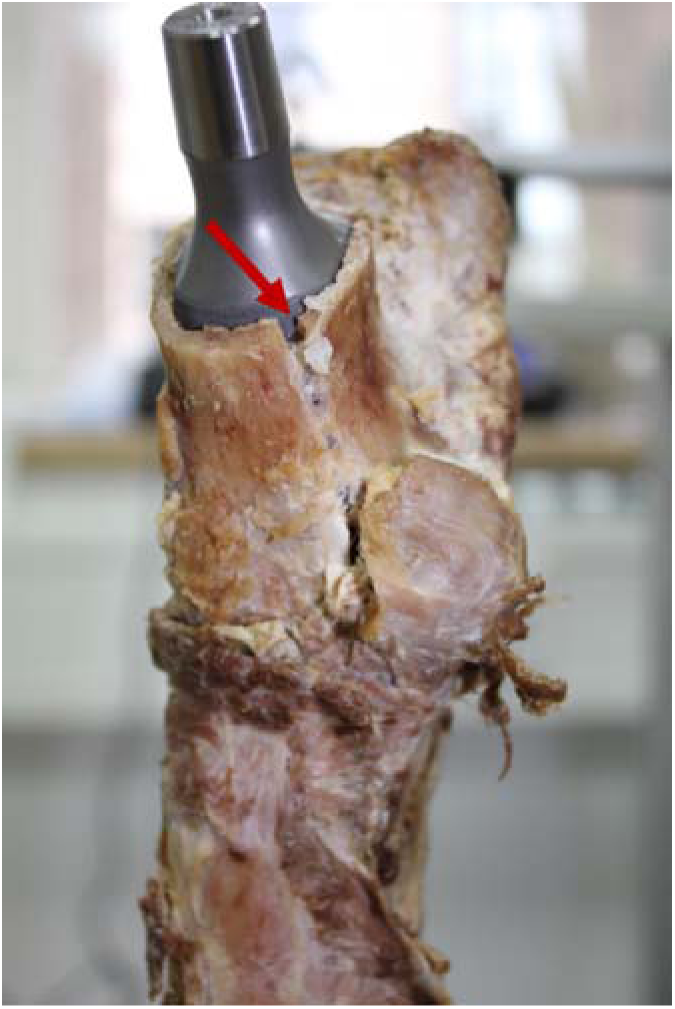
Illustration of a bone fractured during the implantation process.

**Figure 6:**
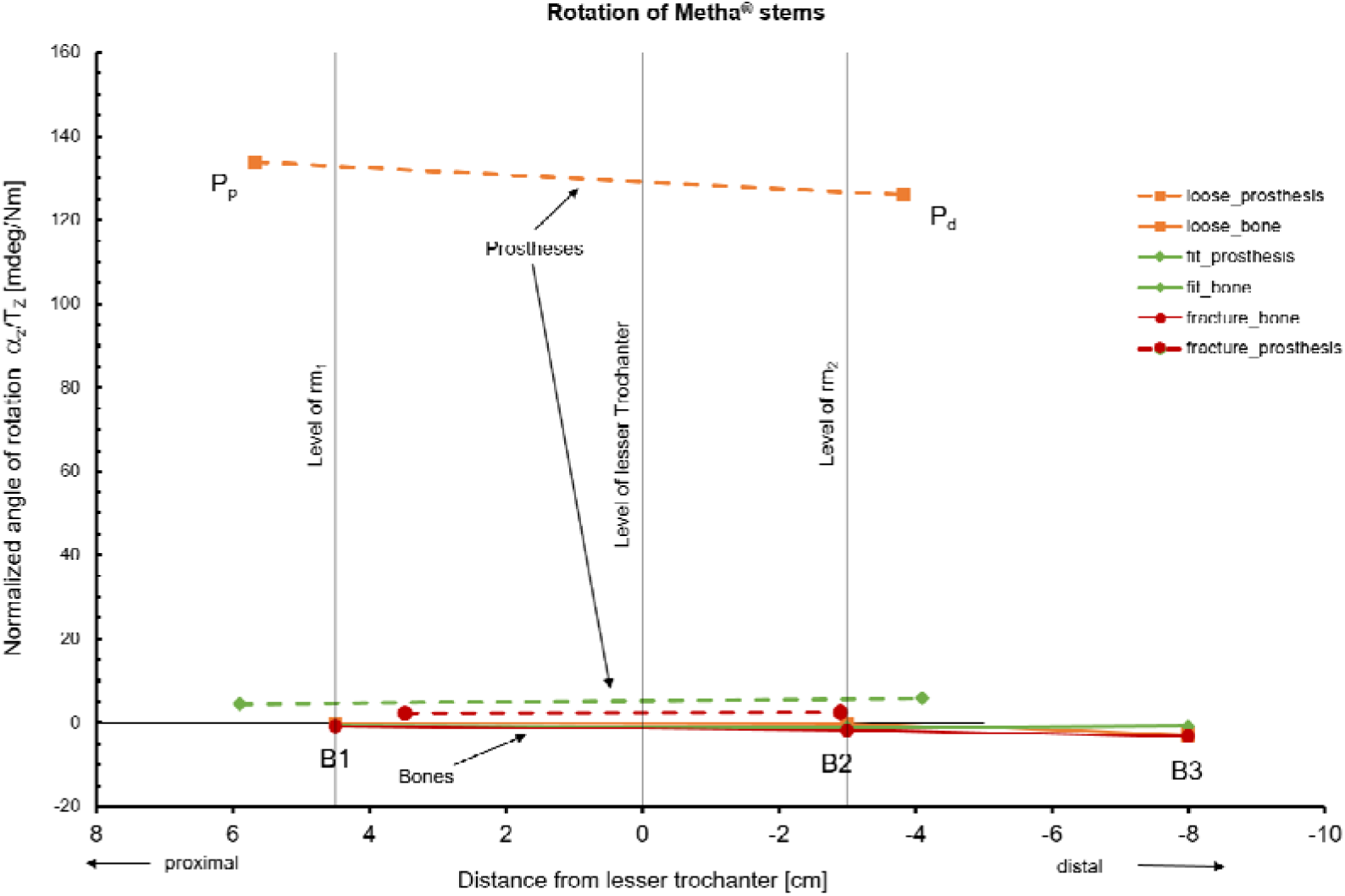
Relative micromovement patterns reveal distinct primary stability states across fixation conditions

### Motion graph

The motion graph illustrates the normalized relative movements between the prosthesis and the femur as a function of the defined measurement heights rm1 (proximal) and rm2 (distal). The respective fixation conditions – loose, fit, and fracture – display characteristic patterns. In the loose group, significantly larger distances between the prosthesis and bone (orange lines) are visible, indicating pronounced micromovements and reflecting a lack of force-locked connection between implant and bone. Movement is markedly pronounced both proximally and distally, suggesting insufficient mechanical stability. In the fit group, the lines are closer together, indicating a stable anchorage between prosthesis and femur and biomechanically confirming good primary fixation. The fracture group shows the smallest distances; the movements are minimal, pointing to an almost rigid connection. However, this rigidity may indicate overpressurization, which is associated with an increased risk of fracture. The graphically depicted distances between prosthesis and bone measurement points thus provide direct insight into the quality of the primary connection and allow for a clear differentiation of stability conditions along the anatomical axis (Figure 6).

### Primary stability analysis

The results of the primary stability analysis revealed clear differences between the three anchorage conditions (*loose*, *fit* and *fracture*), which were statistically evaluated using a non-parametric Kruskal–Wallis test. The *fit* group showed low relative micromotions (rm_1_ = 4.41 ± 2.43 mdeg/Nm; rm_2_ = 5.59 ± 4.77 mdeg/Nm). Even lower values were observed in the *fracture* group (rm_1_ = 1.67 ± 0.60 mdeg/Nm; rm_2_ = 2.72 ± 1.97 mdeg/Nm). In contrast, the *loose* group exhibited significantly higher relative micromotions (rm_1_ = 132.78 ± 171.4 mdeg/Nm; rm_2_ = 124.69 ± 125.07 mdeg/Nm), indicating insufficient mechanical fixation and a potentially increased risk of early loosening. Statistically significant differences were particularly evident between *fit* and *loose* (p = 0.003 for rm_1_) as well as between *fracture* and *loose* (p = 0.006 for rm_2_). A significant difference was also found between *fit* and *fracture* for rm_2_ (p = 0.05; see Table 4).

**Table 4:**
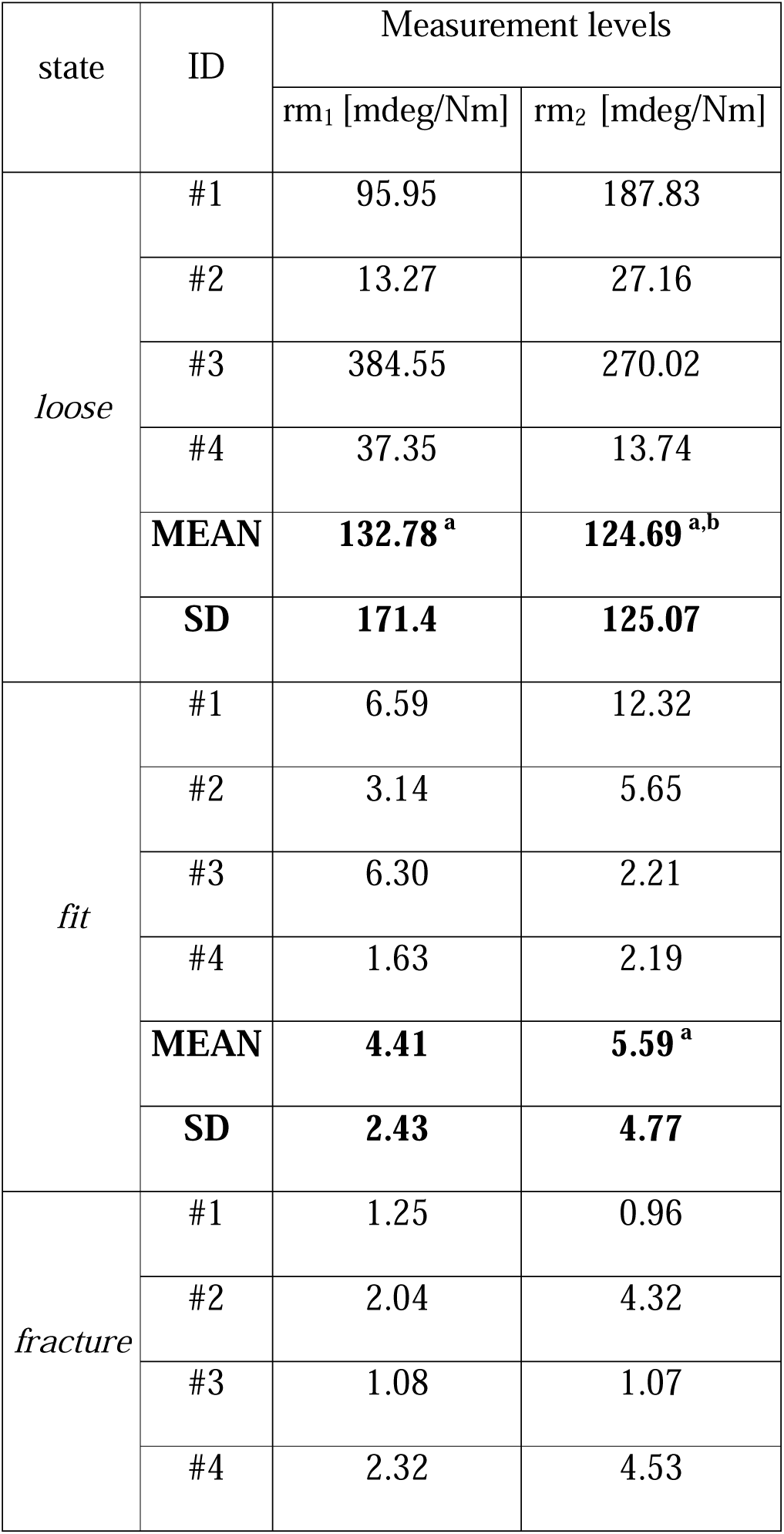
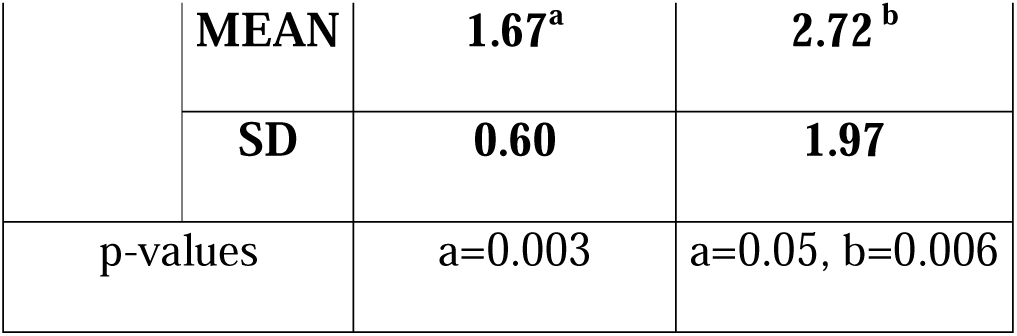
Relative micromotions (rm_1_, rm_2_) distinguish anchorage quality, with significant differences between *loose*, *fit* and *fracture* conditions. Small superscript letters indicate significant differences in pairwise comparison.

## Discussion

The present study aimed to determine whether frequency-domain analysis of impaction sounds can reproduce the spectral markers of primary stability described by Fonseca et al. (Fonseca Ulloa et al. 2023) and, more importantly, whether it can distinguish a clinically hazardous “over-impaction” that culminates in cortical fracture. All hammer blows were recorded under strictly standardized acoustic conditions and processed with an identical FFT-based workflow; statistical inference relied on permutation statistics and family-wise– controlled cluster analysis, thereby avoiding distributional assumptions and α-inflation.

### Re-appearance and shift of the original bands

For the *loose* to *fit* transition the four canonical bands (< 2.5 kHz, 2.9 kHz, 4.4 kHz, 8.7 kHz) were confirmed by cluster permutation, although their effect sizes were small (*d* ≤ 0.14) and global separation minimal (KS = 0.087). The findings are congruent with the original report in which spectral change was attributed primarily to a shift of energy from the low-frequency band to the first two high-frequency bands (Fonseca Ulloa et al. 2023). In the current data this shift is still present but its amplitude is ≤ 0.2 dB, suggesting that the surgeon’s subjective *fit* criterion already limits further seating and thus attenuates the spectral contrast observed in strictly incremental laboratory insertions.

### Emergence of fracture-specific signatures

The transition from *fit* to *fracture* produced a qualitatively different acoustic fingerprint. In the low-frequency window (< 2.5 kHz) band power increased by +6 % (Δµ = 0.062, *d* ≈ 1), whereas energy between 15 and 20 kHz dropped sharply, forming two broad, highly significant clusters (Σ□|*T*| > 5 × 10□). Such high-frequency deficits are characteristic of loss of cortical continuity and the concomitant damping of flexural modes; similar shifts have been reported for microscopic fatigue cracks in cortical bone specimens (Aggelis et al. 2015; García-Vilana et al. 2025; Sakai et al. 2011). The discovery of additional narrow clusters at 7– 9 kHz provides further evidence that the spectral content of the hammer blow is sensitive to small changes in stem-bone coupling long before gross displacement becomes apparent.

### Agreement with mechanical primary-stability testing

Relative micromotion testing demonstrated a clear hierarchy: *fracture* < *fit* < *loose* for both proximal (rm_1_) and distal (rm_2_) relative micromotions. The wide separation between *loose* and *fit* confirms that acoustic changes in the incremental insertion phase mainly reflect global stem seating, whereas the appearance of fracture-specific high-frequency clusters parallels the sudden reduction in micromotion caused by over-press-fit and cortical cracking. The fact that the low-frequency band (< 2.5 kHz) is significantly higher in *fracture* than in *fit* indicates that rigid bony confinement amplifies the fundamental resonance originally observed by Fonseca et al. in the *fit* condition (Fonseca Ulloa et al. 2023). Thus, spectral analysis not only reproduces the classical stability markers but also provides an acoustic “warning zone” above 15 kHz that coincides with the loss of elastic compliance measured mechanically.

### Clinical Implications of Primary Stability

The results underscore the essential importance of primary stability as a key factor for both the short- and long-term success of uncemented short-stem hip prostheses. As the immediate mechanical anchorage of the implant within the surrounding bone, primary stability is crucial for the initial load-bearing capacity of the prosthesis and the subsequent biological integration through osseointegration. Inadequate primary stability can lead to micromovements that impair this integration and significantly increase the risk of aseptic loosening (Bürkner et al. 2012; Hailer et al. 2010).

Biomechanical validation through relative micromotion measurements demonstrated a clear differentiation between anchorage conditions: prostheses in the *loose* group exhibited significantly higher micromotions, indicating a lack of force-locked fixation and thus potentially unstable primary anchorage. In contrast, correctly implanted *fit* prostheses showed minimal motion, confirming sufficient mechanical stability.

Particularly critical is the assessment of the *fracture* group: although relative micromotions were low - superficially suggesting high stability - these cases involved mechanical over-press-fit, resulting in the onset of cortical fractures. Such fractures can easily go unnoticed intraoperatively, especially by less experienced surgeons, as they often do not present immediate clinical or radiographic signs (Adla et al. 2023; Berry 2002). If early-stage fractures remain undetected, there is a considerable risk of complete or displaced fractures during postoperative loading, for instance during early mobilization (Geest et al. 2013). These complications often necessitate complex revision surgeries and significantly delay rehabilitation (Simunovic et al. 2010). Against this backdrop, the combination of acoustic analysis and biomechanical validation offers a valuable tool for intraoperative quality control. The acoustic markers identified in this study provide real-time feedback on the anchorage condition and could serve as an objective early warning system to detect both insufficient fixation and excessive press-fit. This is especially relevant in training situations or in cases with complex anatomy, where the surgeon’s subjective assessment may be limited. In the long term, the results of this study lay the foundation for data-driven decision support in the operating room, with the potential to improve safety during uncemented prosthesis implantation and reduce complications arising from inadequate primary stability.

## Limitations

Despite promising results, this study has several limitations. All tests were conducted on formalin-fixed human cadaveric femora, which differ biomechanically from living bone and lack biological responses. Only one short stem design (Metha®, Aesculap) was examined, limiting generalizability to other implant types. Standardized surgical technique, impaction force, and microphone placement ensured consistency but only partly reflect clinical variability. No direct comparison was made with conventional intraoperative tools (e.g., fluoroscopy), so the diagnostic sensitivity of the acoustic method remains unverified. Lastly, the acoustic markers have not yet been tested in vivo, and their clinical applicability must be assessed in future studies., limiting the ability to compare diagnostic accuracy across modalities.

## Conclusions

This in vitro study demonstrates that spectral cluster analysis of impaction sounds enables objective differentiation between distinct anchorage states of an uncemented short stem hip prostheses. The identified frequency-domain features correlate closely with mechanically measured relative micromotion, thereby providing a reliable acoustic proxy for assessing primary stability.

A particularly noteworthy finding is the detection of high-frequency spectral deficits above 15 kHz combined with a low-frequency gain below 2.5 kHz, which are indicative of incipient cortical fractures. These fracture-specific acoustic signatures not only reflect critical changes in stem-bone coupling but may also serve as early intraoperative warning signals - especially valuable in situations with limited visual control or when procedures are performed by less experienced surgeons.

Overall, the results highlight the technical feasibility and biomechanical validity of real-time acoustic monitoring during prosthesis implantation. This approach offers a promising foundation for developing automated, feedback-supported assistance systems that can enhance intraoperative safety and reduce the incidence of postoperative complications such as loosening or fracture, without disrupting the standard surgical workflow.

## Acknowledgments

### Funding

This work was funded by the Federal Ministry of Education and Research (BMBF, Germany) under grant number FKZ: 16LW0407. The project was managed by the project executing organization VDI/VDE Innovation + Technik GmbH.

### Authoŕs contribution

All authors - A.J., S.S., A.T., S.P., S.H., M.R., and B.I. - were fully involved in the conception, execution, analysis, and interpretation of this study. The authors declare no professional or financial affiliations that could be perceived to have influenced or biased the study’s design, data collection, analysis, or reporting.

